# Generation and first characterization of TRDC-Knockout pigs lacking γδ T cells

**DOI:** 10.1101/2021.03.27.437312

**Authors:** Bjoern Petersen, Robert Kammerer, Antje Frenzel, Petra Hassel, Tung Huy Dau, Roswitha Becker, Angele Breithaupt, Reiner Georg Ulrich, Andrea Lucas-Hahn, Gregor Meyers

## Abstract

The TRDC-Locus encodes the T cell receptor delta constant region, one component of the γδ T cell receptor which is essential for development of γδ T cells. In contrast to peptide recognition by αβ T cells, antigens activating γδ T cells are mostly MHC independent and not well characterized. Therefore, the function of γδ T cells and their contribution to protection against infections is still unclear. Higher numbers of circulating γδ T cells compared to mice, render the pig a suitable animal model to study γδ T cells. Knocking-out the porcine TRDC-locus by intracytoplasmic microinjection and somatic cell nuclear transfer resulted in healthy living γδ T cell deficient offspring. Flow cytometric analysis revealed that TRDC-KO pigs lack γδ T cells in peripheral blood mononuclear cells (PBMC) and spleen cells. The composition of the remaining leucocyte subpopulations was not affected by the depletion of γδ T cells. Genome-wide transcriptome analyses in PBMC revealed a pattern of changes reflecting the impairment of known or expected γδ T cell dependent pathways. Histopathology did not reveal developmental abnormalities of secondary lymphoid tissues. However, in a vaccination experiment the KO pigs stayed healthy but had a significantly lower neutralizing antibody titer as the syngenic controls.

## Introduction

The adaptive immune system is composed of three lymphocyte subsets, B cells, T cells expressing the αβ T cell receptor (TCR) and T cells expressing γδ TCRs. The tripartite organization of the lymphocytic immune system seems to be fundamental since it evolved independently in jawed and jawless vertebrates^1^. These cells generate their antigen specific receptors using somatic V(D)J recombination, which enables them to recognize a vast spectrum of different antigens. While the nature of antigen recognition by B cells and αβ T cells is well established, a unifying concept for γδ T cell antigen recognition and function is still pending. The majority of γδ T cells are activated in an MHC-independent manner, which is in striking contrast to MHC-restricted αβ T cells. They can attack target cells directly through their cytotoxic activity or indirectly through activation of other immune cells^2^. Until recently, γδ T cells were thought to be simply innate immune cells with limited or redundant functions^3^. The current view is that these cells complement many different players of the immune system^4^, and it is becoming obvious that they represent a heterogenous population of cells with important unique features in many infections, autoimmune diseases, allergies and in immunoregulation. A wide range of γδ T cell functions have been described in humans and mice, including skin and mucosal epithelial wound repair, induction of tolerance, cytotoxicity and the production of various cytokines that regulate immune responses ^5–8^. The increase in γδ T cells following vaccinia virus infection, high vaccinia virus replication in γδ T cell knockout mice, and the selective lysis of γδ T cells against vaccinia virus infected target cells indicate some role for γδ T cells during virus infections^9–11^. To understand what they do and what role they play in different challenging situations of the immune system, a knockout animal model would be indispensable. A γδ T cell deficient mouse model was successfully established^12^, however γδ T cells show a remarkable degree of diversification in function, anatomical localization and TCR usage that also differs largely between species^13^. Therefore, care has to be taken when extrapolating results from one species to the other. In humans and mice γδ T cells constitute only 0.5–10% of T cells in peripheral blood but are substantially enriched in epithelial tissues (e.g. in skin, lungs, intestine). A prominent population of γδ T cells in human blood, V*γ*9V*δ*2 T cells which recognize phosphoantigens (PAgs) do not exist in mice but were recently described in the new world camelid *Vicugna pacos* (alpaca) ^14^. Furthermore, while in human and mice a limited number of γδ VDJ cassettes exist, species like cattle, pigs and chicken have larger numbers of γδ VDJ cassettes and a high percentage of circulating γδ T cells (20 - 50 %), thus called γδ T cell-high species in contrast to γδ T cell-low species (humans and mice)^15^. In addition, in some species the co-receptor WC1 family, which belong to the group B scavenger receptor cysteine-rich molecules, is expanded ^16^. WC1 receptors are exclusively expressed by γδ T cells and function as hybrid pattern recognition receptors and γδ TCR co-receptors in cattle ^17^. Bovine WC1 receptors can directly bind to pathogenic bacteria and transduce activation signals to γδ T cells, thus, γδ T cells may be crucial for the control of certain systemic infections ^17^. Based on the remarkable lack of conservation between γδ T cells of different species, additional animal models are needed to understand the function of γδ T cells and to establish a unifying concept for their role during immune responses. In this report we describe the generation of TRDC-KO pigs and their genetically identical wild type controls. Since the number and quality of inbred pig strains are limited, the identity at the MHC locus of the KO and control animals is of utmost importance for the comparison of the immune response to infections. We did not observe any abnormalities in health and behavior of the KO pigs under standard housing conditions. In addition, the immune system in KO pigs developed normally, with the exception of the absence of γδ T cells. The availability of a γδ T cells deficient animal model from a γδ T cell-high species may add important new insights in the role of this mysterious part of the adaptive immune system.

## Results

### Intracytoplasmic microinjection of porcine *in vitro* derived zygotes with TRDC-CRISPR/Cas and transfer to recipients

After transfer of 35 microinjected embryos, both recipients (#695, #696) were determined pregnant on day 25 after surgical embryo transfer. Sow 695 delivered 5 liveborn piglets on day 112 of gestation and sow 696 gave birth to 9 liveborn piglets on day 114 of gestation (Table 1). All piglets were healthy and showed no aberrations regarding birth weight compared to age-matched counterparts or wild type littermates.

**Table 1:**
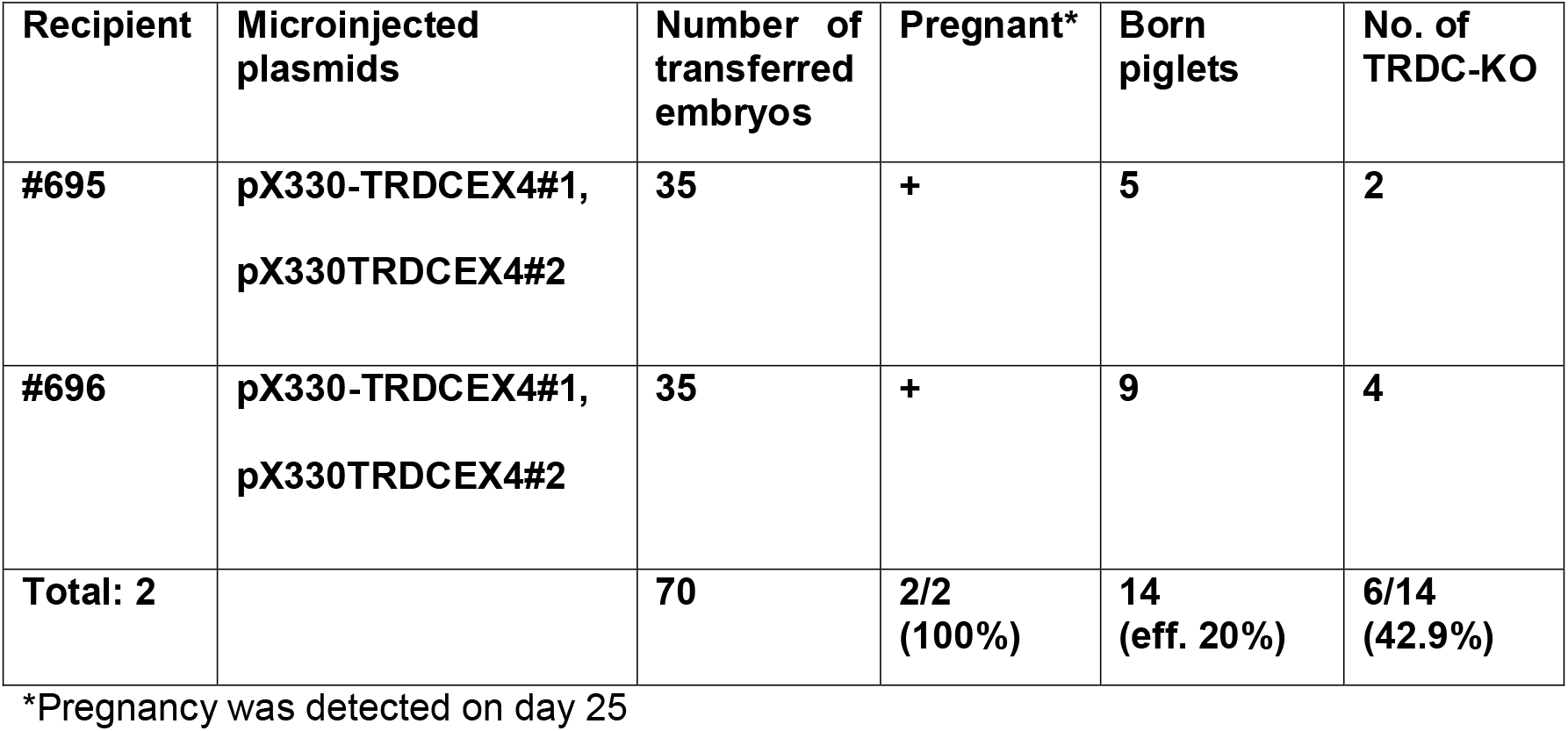
Results of the intracytoplasmic microinjection of CRISPR/Cas9 targeting exon 4 of the porcine TRDC locus in porcine zygotes.

| Recipient | Microinjected plasmids | Number of transferred embryos | Pregnant* | Born piglets | No. of TRDC-KO |
| --- | --- | --- | --- | --- | --- |
| #695 | pX330-TRDCEX4#1,<br>pX330TRDCEX4#2 | 35 | + | 5 | 2 |
| #696 | pX330-TRDCEX4#1,<br>pX330TRDCEX4#2 | 35 | + | 9 | 4 |
| **Total: 2** |  | **70** | **2/2**<br>**(100%)** | **14**<br>**(eff. 20%)** | **6/14**<br>**(42.9%)** |

\*Pregnancy was detected on day 25

### Genotyping of TRDC-KO piglets

Genomic DNA from tail tips was used for PCR-based detection of genetic modifications at the TRDC locus in the piglets. The genomic DNA of the piglets originating from intracytoplasmic microinjection of the pX330-TRDCEX4 #1 and #2 plasmids was employed for PCR (Figure 1) and subsequent sequencing (Figure 2) to detect genetic modifications at the TRDC gene. In total, two out of five piglets (40%) from sow 695 showed a biallelic 40 bp deletion within exon 4 of the TRDC gene, while four out of nine piglets (44.4%) from sow 696 carried a biallelic 40 bp deletion. The remaining piglets were detected to be wild type. The overall efficiency to generate a 40 bp biallelic deletion in the exon 4 of the TRDC gene was 42.9% (6/14).

**Figure 1:**
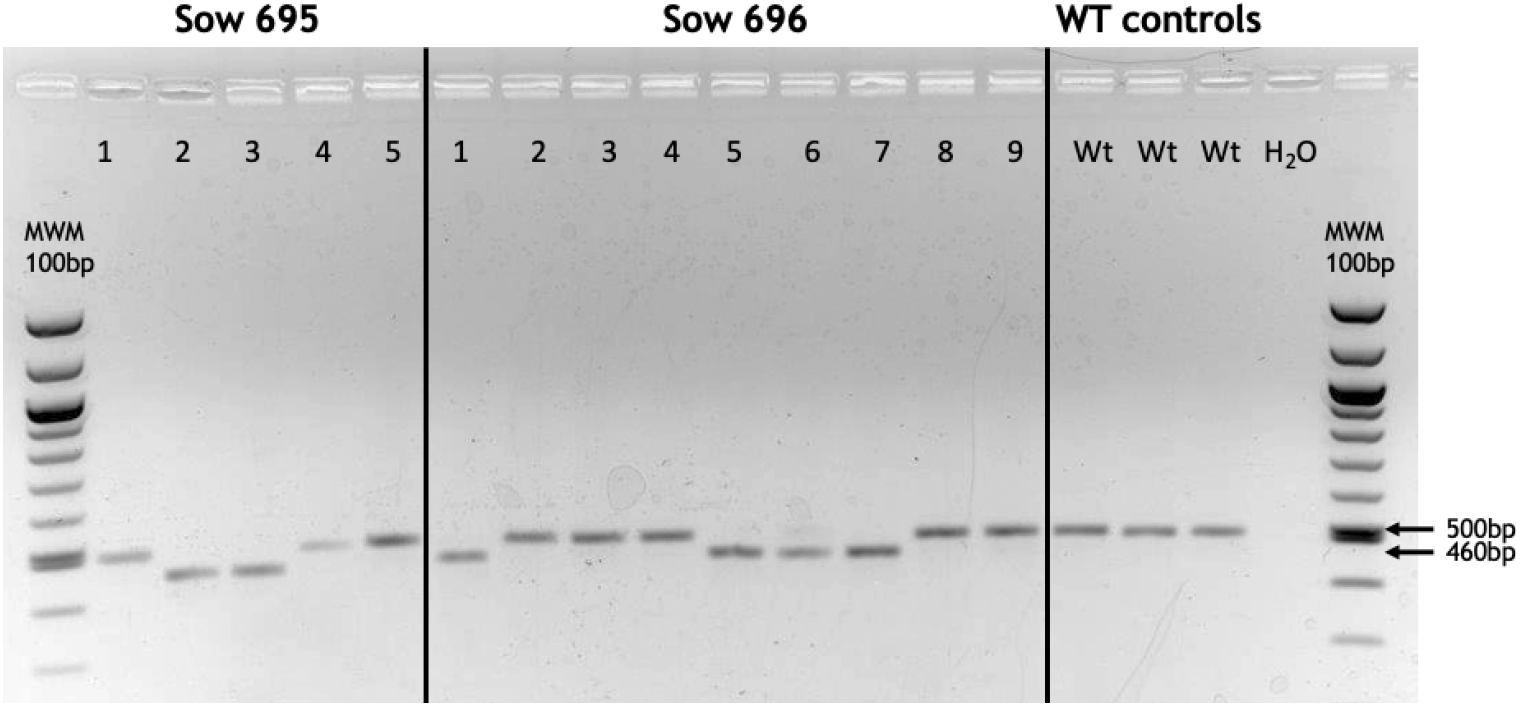
PCR-Analysis of the porcine TRDC locus in samples from pigs generated by intracytoplasmic microinjection. Pigs 695-2, −3 and 696-1, −5, −6, −7 display a lower band of about 460 bp in contrast to wild type pigs (500 bp) indicating a 40 bp deletion caused application of the CRISPR/Cas9 construct.

**Figure 2:**
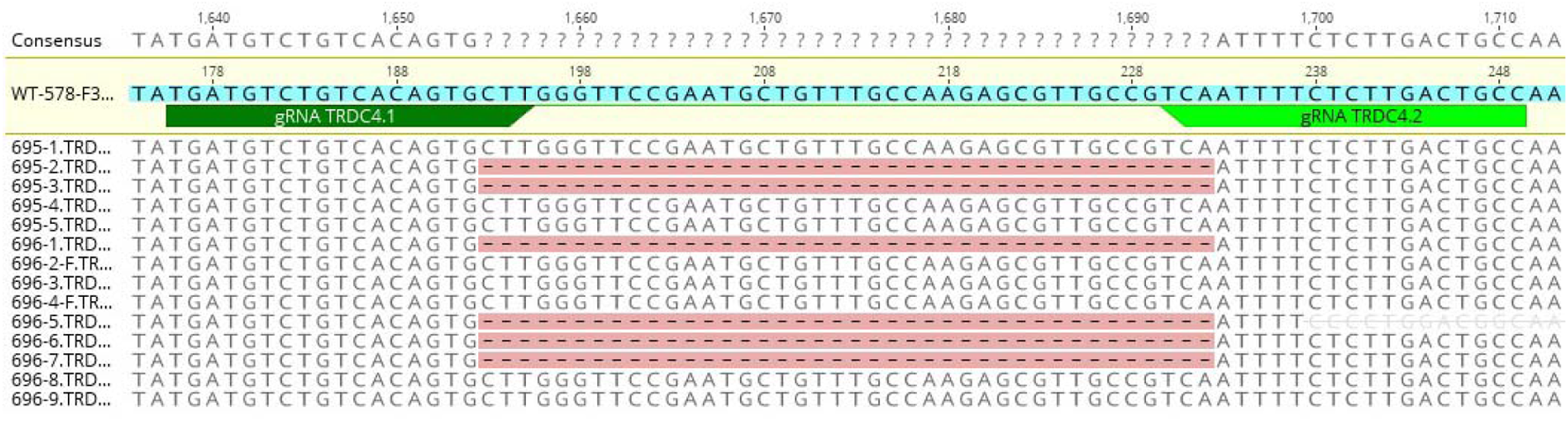
Sequencing data of pigs generated by intracytoplasmic microinjection of the CRISPR/Cas9 construct targeting the porcine TRDC-locus. Sequencing results confirmed the PCR results (Fig.1) that pigs 695-2, −3 and pigs 696-1, −5, −6, −7 carry a homozygous knockout of the TRDC locus consisting of a clear 40 bp deletion. For each piglet, six samples were sequenced. None of the piglets were monoallelic knockouts. (The binding sequences of the guide RNAs are indicated by green arrows).

### Transfection of CRISPR/Cas9 plasmids

As an alternative to the intracytoplasmic microinjection approach, we employed somatic cell nuclear transfer to produce genetically identical TRDC knockout pigs and syngeneic wild type control pigs. Therefore, 3×10^6^ cells in total were co-transfected with the plasmids pX330-TRDCEX4#1 and #2 and subsequently seeded on three T25 flasks (25 cm^2^). After reaching 70-80 % confluency, cells were detached by EDTA/Trypsin treatment and seeded at a concentration of 7-10 cells per well on a 96-well plate. Individual cell clones were analyzed by PCR. Cells with a biallelic deletion of the TRDC gene (−40 bp) showed only the lower band, while cell cultures with a mixed population of wild type, monoallelic and biallelic genetically modified cells showed an upper wild type band at 499 bp and a lower band at 459 bp (Figure 3). Cell clones D12 (mixture) and H2 (almost pure lower band) were chosen as donor cells for SCNT. D12 was chosen to produce both TRDC knockout and syngeneic wild type control pigs.

**Figure 3:**
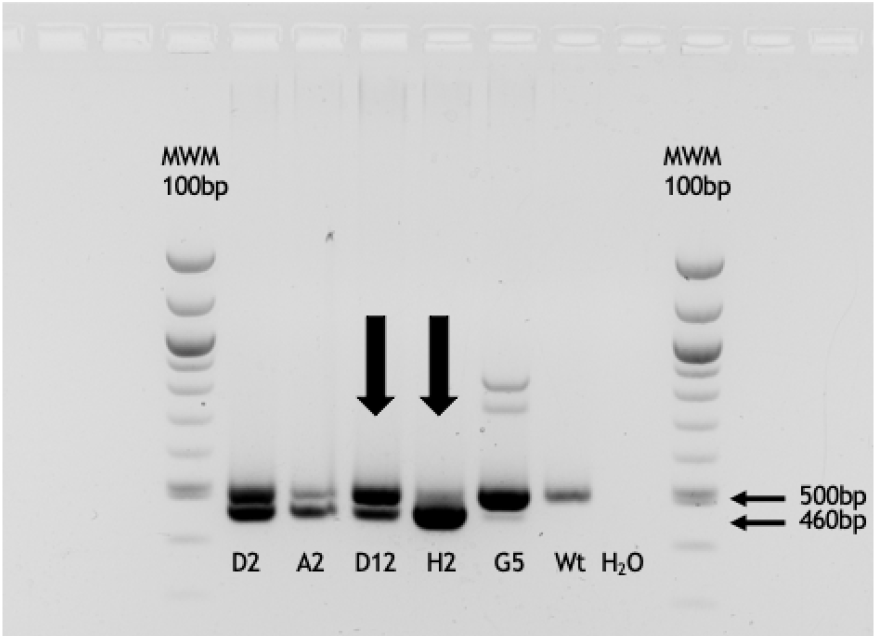
PCR analysis of the TRDC locus of CRISPR/Cas9 transfected cells. The lower band of 460bp indicates a 40bp deletion caused by the CRISPR/Cas9 construct. Cell clone D12 displayed two bands of 500 and 460bp of size which indicates a mixed population of cells consisting of TRDC-knockout cells, wild type cells and/or cells with a monoallelic knockout of the TRDC locus. In contrast, cell clone H2 shows a single band of 460 bp indicating an almost pure population of TRDC knockout cells.

### Somatic cell nuclear transfer of TRDC-KO embryos and syngeneic controls

H2-derived cloned embryos were surgically transferred to two hormonally synchronized recipients. Each of the two recipients (#8106 and #738) received 90 cloned embryos and were detected to be pregnant on day 25 after embryo transfer. D12-derived cloned embryos were also transferred to two recipients. One recipient (#8115) received 97 embryos, while the second one received 96 embryos (#737). Again, both recipients were determined to be pregnant by ultrasound scanning on day 25 after embryo transfer (Table 2). All recipients were allowed to go to term and delivered in total 9 healthy liveborn (9/17:52.9%) and 8 stillborn piglets (8/17:47.1%). The stillborn piglets did not show any abnormalities but were not checked for deletions in the exon 4 of the TRDC gene.

**Table 2:** Results from somatic cell nuclear transfer employing genetically modified cells as donor cells.

| Recipient | Cell clone* | Number of transferred embryos | Pregnant | Born piglets (stillborn) | No. of TRDC-KO of liveborn |
| --- | --- | --- | --- | --- | --- |
| #8106 | H2 | 90 | + | 2 (2) | n.d. |
| #738 | H2 | 90 | + | 3 (1) | 2/2 |
| #8115 | D12 | 97 | + | 3 (3) | n.d. |
| #737 | D12 | 96 | + | 9 (2) | 4/7 |
| **Total: 4** |  | **373 (avg. 93.3)** | **4/4 (100%)** | **17 (cloning efficiency 4.6%)** | **6/9 (66.7%)** |

\* cell clone D12: mixed population of knockout and wild type cells, cell clone H2: pure knockout cells.

PCR analysis of the cloned piglets revealed that 4 out of the 7 piglets (57.1%) originating from the mixed cell clone D12 (737-1-7) carried a biallelic 40 bp deletion of the TRDC gene, while the remaining 3 piglets remained wild type (42.9%). The two piglets (738-1, −2) originating from cell clone H2 carried a biallelic 40 bp deletion of the TRDC gene, as expected (Figure 4).

**Figure 4:**
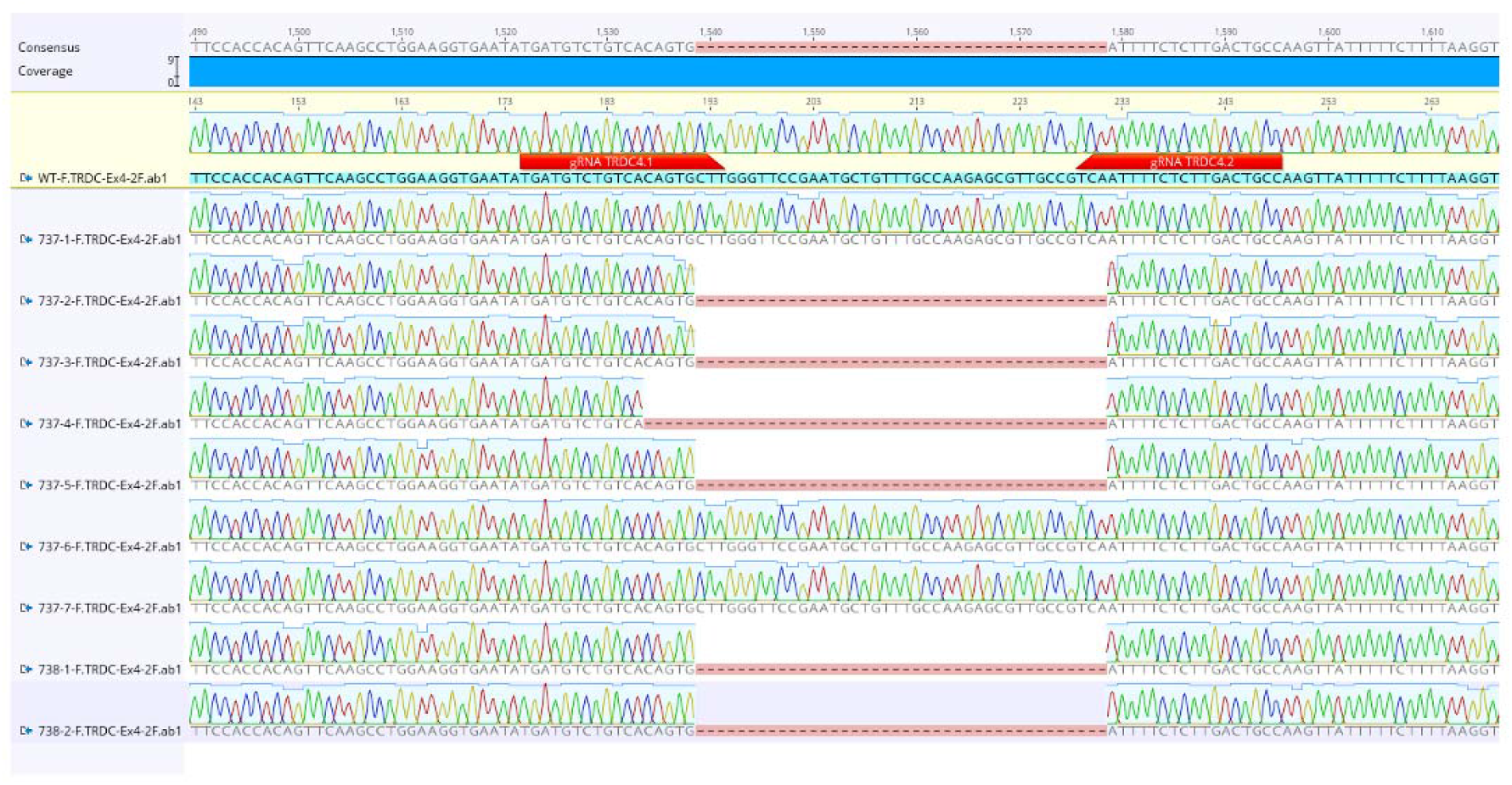
Sequencing results of the cloned liveborn TRDC knockout pigs. Pig 737-4 differed regarding the genetic modification compared to the other TRDC knockout pigs with a larger deletion encompassing 45bp, indicating its origin from a separate cell subclone. The binding sequences of the guide RNAs are indicated by red arrows.

### Fitness and viability of TRDC-knockout pigs

The TRDC knockout pigs did not differ to syngenic wild type control animals and age-matched wild type pigs in our facility in regard of their health status and growing performance (Figure 5). TRDC-KO pigs are kept for 2 years in our facility under standard conditions and never showed any health impairment.

**Figure 5:**
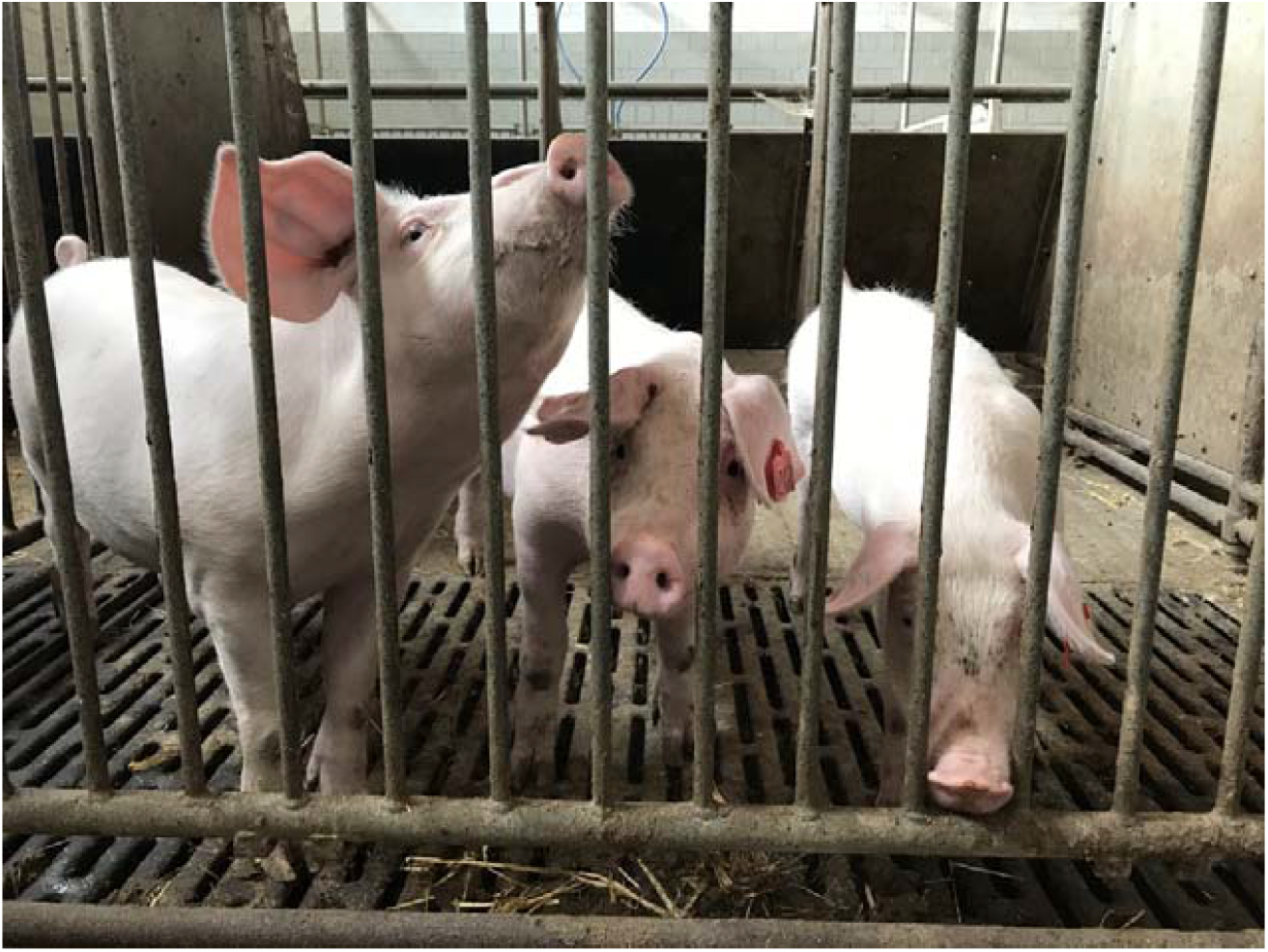
Three TRDC-knockout pigs from intracytoplasmic microinjection.

### FACS analysis of TRDC-knockout blood and spleen cells

To determine the effect of the 40 bp deletion in the TRDC gene on lymphocyte composition in TRDC-KO pigs we performed flow cytometry analysis of PBMC and spleen cells from 8 wt pigs and 5 TRDC-KO pigs. We used two mAb that specifically detect independent antigens of porcine γδ T cells. One binds to the δ-chain of the T cell receptor and the other to the CD3 antigen of γδ T cells. In average 20 % (ranging from 8 % – 47 %) of peripheral blood T cells were double-stained by these mAbs in wild type pigs while no double-stained cells were detected in TRDC-KO pigs, indicating that no γδ T cells were present in peripheral blood of TRDC-KO pigs (Figure 6A). In addition, we isolated spleen cells from wild type and TRDC-KO pigs and analyzed the content of γδ T cell populations. Typically, in wild type pigs the majority (80 %) of γδ T cells in the blood belong to the CD2 negative phenotype, while in the spleen the majority (66 %) of γδ T cells are CD2 positive. No γδCD3 positive cells of either type were detected in the spleen of TRDC-KO pigs (Figure 6B). These data indicate that no γδ T cells developed in TRCD-KO pigs. The majority of γδ T cells in pigs are CD4 and CD8 negative, reflected by minimal numbers of CD4/CD8 double negative T cells in TRCD-KO pigs (Figure 6C). Comparing the numbers of various lymphocyte subpopulations between TRDC-KO pigs and wild type pigs we did not find significant differences (Figure 6D). These results indicated that the absence of γδ T cells had no major impact on the development of other lymphocyte populations.

**Figure 6:**
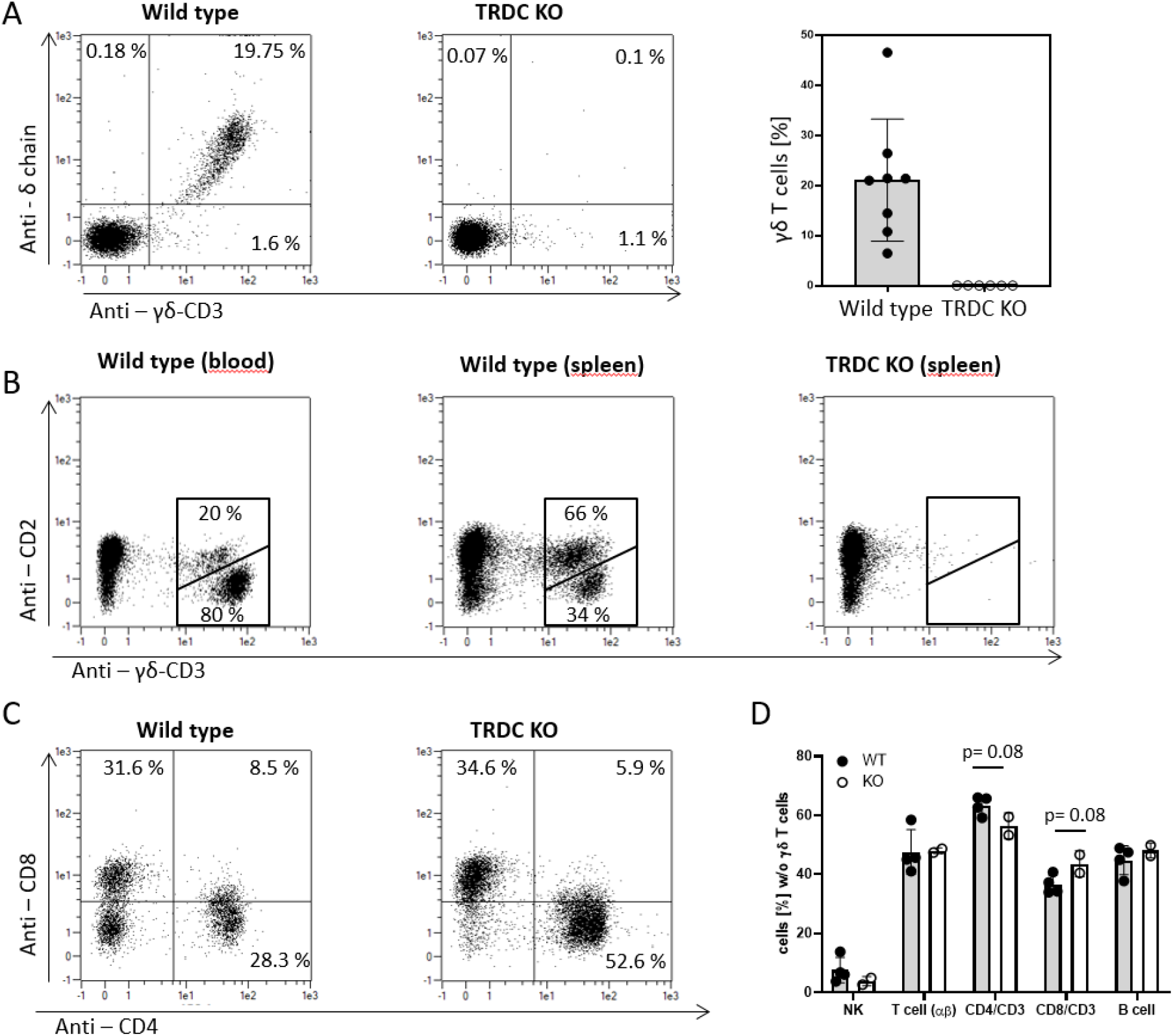
No γδ T cells are present in blood and spleen of TRDC-Knockout pigs. **(A)** PBMC of TRDC-Knockout and wild type pigs were stained with anti-CD3, anti-δ chain and anti-γδ T cell-specific CD3 mABs. CD3 positive cells were gated and δ chain vs. γδ T cell-specific CD3 dotblots are displayed. About 20% of peripheral blood T cells were γδ T cells in wild type pigs while no γδ T cells were detected in TRDC-Knockout pigs. **(B)** PBMC and spleen cells were isolated and staind for CD2 and γδ T cell-specific CD3. Typically in wild type pigs the majority (80%) of γδ T cells in the blood belong to the CD2 negative phenotype (left dot blot), while in the spleen the majority (66%) of γδ T cells are CD2 positive (middle dot blot). No γδ T cells of either type were detected in the spleen of TRDC-Knockout pigs. **(c)** Most γδ T cells are CD4 and CD8 negative. Accordingly TRDC-Knockout pigs have minimal numbers of CD4/CD8 double negative T cells. **(D)** The percentage of lymphocyte subpopulations of all lymphocytes without γδ T cells were compared of wild type and TRDC-Knockout pigs. No significant difference of any lymphocyte subpopulation was detected.

### Comparison of gene expression profiles of TRDC-KO versus control animals via microarray analysis

Since we did not detect a major influence on the composition of lymphocyte subpopulations in the peripheral blood by the complete loss of γδ T cells, we asked whether the absence of γδ T cells has an impact on the transcriptional profile of the remaining lymphocyte population. Thus, we performed microarray analysis of PBMC isolated from wild type pigs and TRDC-KO pigs. Only 0.7 % of genes were regulated more than 2-fold (0.4 % upregulated and 0.3 % downregulated) in PBMC of TRDC-KO pigs (Figure 7A). Hierarchical clustering identified gene clusters with a significant expression level in either the wild type or the TRDC-KO pigs, only these genes were further analyzed (Figure 7B). A high portion of down-regulated genes are known to be preferentially expressed by γδ T cells, such as δ T cell receptor genes, GATA3, WC1.1, SOX13, IL-1R and IL-18R. Genes which were upregulated in TRDC-KO pigs encompass genes which are preferentially expressed by granulocytes and monocytes in human PBMC like Zyxin, CXCL8 and CXCL2. Thus, the transcriptional profile of PBMC in TRDC-KO pigs reflect the loss of γδ T cells and as a result the proportional increase of non-lymphoid cells.

**Figure 7:**
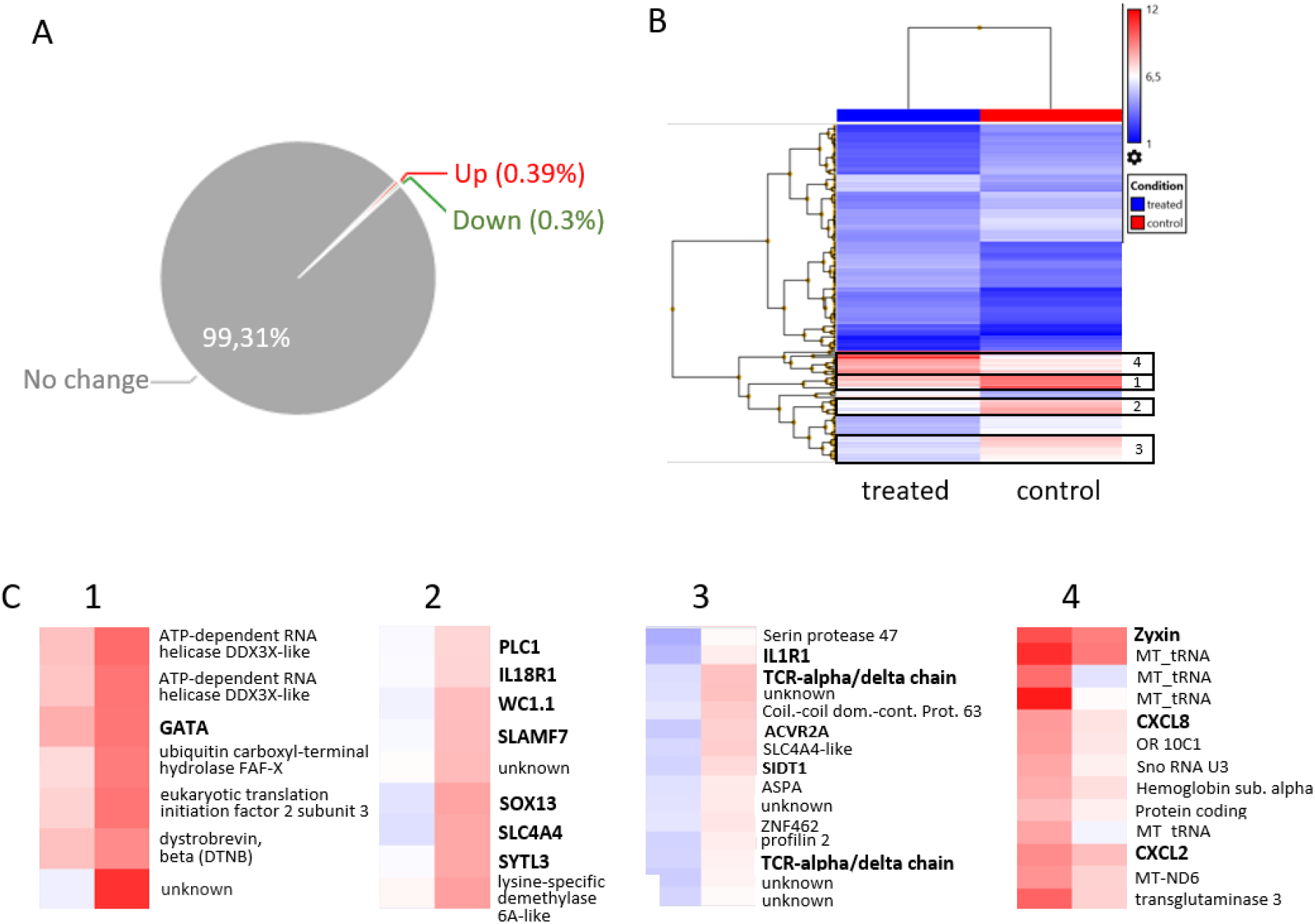
Transcriptional changes in PBMC of TRDC-Knockout pig refelect the loss of γδ T cells. **(A)** Only 0.7 % of genes were regulated more than 2 fold in PBMC of TRDC-Knockout pigs. **(B)** Using hirachical clustering up-regulated and down-regulated genes were sorted acording to the expression levels. **(C)** down-regulated (1,2,3) and upregulated (4) genes from the hirachical clustering (B) were characterized in more detail. (1) all annotated genes of this cluster were expressed preferentialy by T cells. GATA3 is highly expressed by γδ T cells. (2) Cluster 2 contains genes like WC1.1 and SOX13 which are preferentially expressed by γδ T cells to a various degree. (3) T cell receptor genes were found in cluster 3. In addition, IL-1R (3) and IL-18R (2) were downregulated in PBMC of TRDC-Knockout pigs. (4) Genes which were upregulated contain genes which were preferentially expressed by granulocytes and monocytes in human PBMC like Zyxin, CXCL8 and CXCL2.

### No differences between wild type and TRDC-KO pigs were detected by gross pathological, histological and immunohistological examination of lymphoid tissues

We wondered if the development of lymphoid tissues was influenced by the absence of γδ T cells. At autopsy the size and morphology of all lymphoid tissues did not differ between wild-type or knockout pigs. Further, neither the detailed histopathologic evaluation on H&E stained slides nor the application of T-cell (CD3) and B-cell (CD20) markers revealed differences in the overall architecture, distribution and quantity of the immune cells labelled. Representative slides of the thymus, ileal Peyer’s patches and tonsil are shown in Figure 8 (A-R). Both mAbs directed against the specific γδ T cell-antigens (PGBL22A, PPT16) cross react on tissue slides with cytoplasmic structures of an unknown cell type, resulting in unspecific reactions. Thus both antibodies we had at hand, were unfortunately not suited for the use in immunohistology on cryosections.

**Figure 8:**
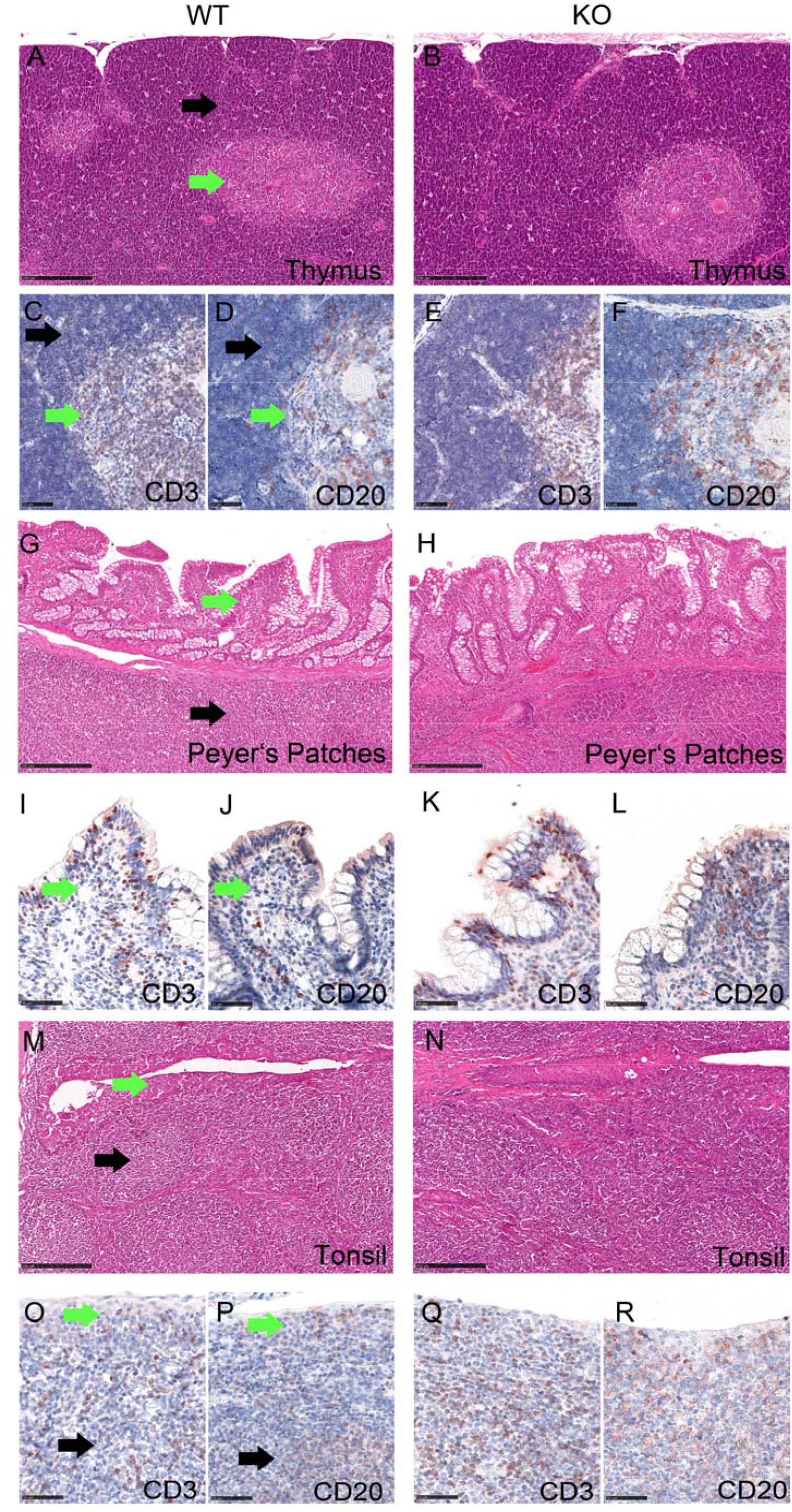
No differences between wildtype and TRDC-Knockout pigs were detected by histological examination of lymphoid tissues. Lymphoid tissues from wild type (WT: A, C, D, G, I, J, M, O, P) and knockout (KO: B, E, F, H, K, L, N, Q, R) domestic pigs. Thymus (A, B) with cortex (black arrow) and medulla (green arrow), Peyer’s Patches (G, H) with mucosa associated lymphoid follicle (black arrow) and absorptive epithelium (green arrow) containing intraepithelial, and transmigrating lymphocytes, Tonsil (M, N) with mucosa associated lymphoid follicle (black arrow) and mucosal epithelium (green arrow) containing intraepithelial and transmigrating lymphocytes. H&E stain (A, B, G, H, M, N), immunohistochemistry (C-F, I-L, O-R), using anti-CD3 or anti-CD20 antibody as indicated, ABC method, AEC chromogen

### Possible role of γδ T cells in vaccine-mediated protection

As a first approach towards testing the immunological competence of the TRDC-KO pigs we decided to vaccinate the animals with a commercial vaccine against the pestivirus classical swine fever virus (CSFV). The vaccine contains a highly attenuated live virus of the so-called C-strain type. In normal pigs, the vaccine is known to be completely apathogenic but very effective with regard to induction of a protective immunity against CSFV field virus infection^18–21^. Three knockout pigs [group 1, numbers 1314, 1318, 1319] and two syngenic wild type controls [group 2, numbers 1315, 1320] were vaccinated at day 0 with the commercial live virus vaccine via the intramuscular route as recommended by the supplier. Rectal temperature of the animals was recorded daily starting 10 days before vaccination. The animals were monitored daily for general health status. Neither in group 1 nor 2 any signs of disease were detected during the observation period. Body temperatures of all animals remained in the normal range. Blood for serum production was taken on day 0 and on day 21 p.i. The sera were tested for the presence of neutralizing antibodies against the homologous CSFV C-strain. As expected, the results were negative for all animals at day 0, proving that they had not had any contact to CSFV antigens before. In contrast, neutralization titers were recorded on day 21 for all animals (Figure 9). The values were significantly higher for the wild type pigs compared to TRDC-KO (P value = 0.0428).

**Figure 9:**
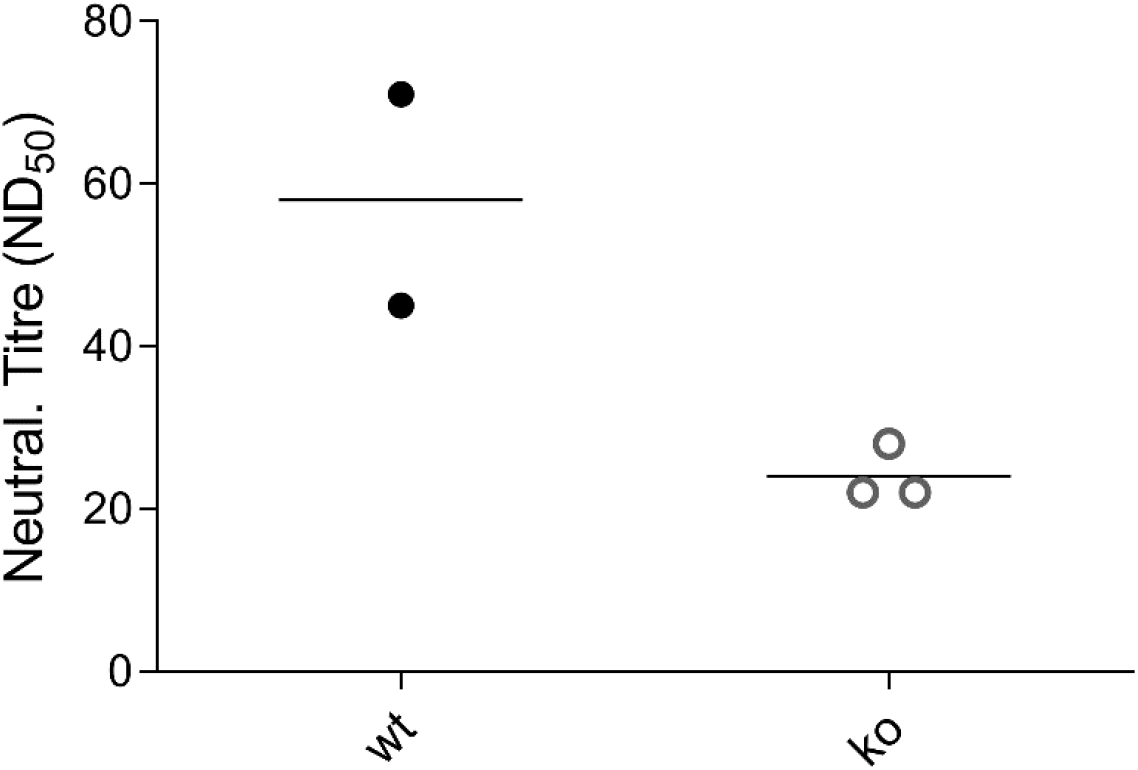
Diagram showing the titers of virus-neutralizing serum antibodies of 2 syngenic wt (wt) and 3 TRDC-Knockout (ko) pigs at day 21 post vaccination, given as the maximal reciprocal dilution of the sera that is able to neutralize 50% of 100 TCID_50_ infective doses of CSFV C-strain calculated according to Spaerman-Kaerber. The mean values for both groups are shown as horizontal lines. The P value determined by statistical analysis is 0.0428.

## Discussion

Mice deficient for γδ T cells have been established almost 30 years ago^22^. In the meantime, they have proven to be extraordinary helpful to elucidate functions of γδ T cells. In particular, the role and contribution of certain murine γδ T cell compartments during infections with viral, bacterial and parasitic pathogens could be defined using these mice^23–25^. Taken into account that γδ T cell populations can differ largely between species it is of considerable importance to have γδ T deficient animals from additional species. One major variation of γδ T cell compartments between species is the amount of circulating γδ T cells. The proportion of circulating γδ T cells is surprisingly high in artiodactyls (e.g. cattle, sheep and pigs) and chicken^15^. Thus, the animal model described in the present report is the first of a γδ T cell-high species. Pigs with a biallelic knockout of the TRDC gene were efficiently produced either by intracytoplasmic microinjection or somatic cell nuclear transfer using modified cells as donor cells. While healthy offspring could be obtained after embryo transfer of microinjected zygotes, somatic cell nuclear transfer (SCNT) resulted in 47.1 % stillborn piglets. As healthy TRDC knockout piglets were obtained by both techniques, we conclude that the relatively high number of stillborn piglets was associated with SCNT and false or incomplete reprogramming of the donor cell nucleus ^26–28^. TRDC knockout pigs developed normally and reached sexual maturity. After breeding TRDC deficient pigs, all offspring carried a monoallelic knockout of the TRDC gene as expected. Though mosaicism cannot be excluded in the pigs originating from intracytoplasmic microinjection^29^, we found the 40 bp deletion in all tissues analyzed (skin, heart, liver, kidney, muscle), rendering mosaicism unlikely. Besides, no off-target events could be detected in the 10 most possible genomic sites in microinjected and cloned offspring (data not shown). As previously observed in Tcrd−/− mice no γδ T cells remain in the TRDC-KO pigs while the studies suggest the presence of a normal αβ T cell repertoire in the periphery of the γδ T cell deficient pigs^22^. Furthermore, microarray analysis did not indicate that there is a major impact on the transcriptome of PBMC by the loss of γδ T cells. These observations may explain why the immune phenotype of TRDC-KO pigs is surprisingly mild. The vaccination experiment described above was conducted with a commercial vaccine containing a live virus that is known to be completely apathogenic in wild type pigs ^18–21^. We have chosen this approach to check whether the knockout pigs are hampered in a way that they cannot control a virus infection at all, even when the infecting virus is highly attenuated. In fact, we saw no signs of disease in the vaccinated pigs which could be explained in different ways, namely that the so far unknown attenuation principle of CSFV C-strain is sufficient even in immunologically compromised animals, that the loss of γδ T cells has only mild effects on virus control by the immune system or a mixture of both reasons. The results of the antibody neutralization test showed that the knockout pigs were able to mount a measurable antibody response to an infecting virus, but the titers were lower than in the wild type animals. Even though the statistical significance of this result is only low due to the small number of animals tested, it represents a first indication for a reduced immunological competence of the knockout animals. This may indicate that different outcomes of infections are due to the missing contribution of γδ T cells during the immune response not to other deficiencies of the immune system. This point will have to be followed up using different infection and challenge systems. There are various reports pointing to a protective role of γδ T cells against important pig pathogens such as African swine fever virus^30,31^, Foot and mouth disease virus^32,33^, Mycobacterium bovis^34^, Porcine respiratory and reproductive syndrome virus^35,36^, and Influenza A virus ^37^. A specific role of γδ T cells has not been definitively proven so far, but with the TRDC-KO pigs at hand, the large variety of swine pathogens displaying different ways of pathogenesis and interplay with the hosts immune system can be tested to finally elucidate the role of γδ T cells in immunological defense against infectious agents. The fact that some of these pathogens are zoonotic and responsible for severe human diseases render this approach even more attractive.

## Material and Methods

### Animals

German Landrace pigs served as recipient animals for genetically modified embryos derived by somatic cell nuclear transfer (SCNT).

### CRISPR/Cas vector and single-guide RNA

The CRISPR/Cas9 system was used to induce defined deletions within the exon 4 of the porcine TRDC gene (Ensembl transcript: ENSSSCT00000026772). Guide RNAs (gRNAs) were designed using the web-based design tool *CRISPOR* (http://crispor.tefor.net/) (Fig. X). Target sequences were analyzed via BLAST to reduce the probability for off-target events. Two oligo duplexes including the target sequence (TRDCEX4#1: 5’-CAC CGT GAT GTC TGT CAC AGT GCT T-3’ and TRDCEX4#2: 5’-CAC CGG CAG TCA AGA GAA AAT TGA-3’) and BbsI overhangs were designed, and each gRNA was cloned into a linearized CRISPR/Cas9 vector pX330-U6-Chimeric_BB-CBh-hSpCas9 (Addgene plasmid # 42230)^38^. The final plasmids pX330-TRDCEX4#1and pX330-TRDCEX4#2 were then used for intracytoplasmic microinjection into IVF zygotes and for transfecting porcine fetal fibroblasts.

### Culture and lysis of primary cell cultures from wild type fetuses

Porcine fibroblasts were isolated from ear tissue of wild type piglets and cultured in Dulbecco’s modified Eagle’s medium (DMEM) with 2 % penicillin/streptomycin, 1 % non-essential amino acids and sodium pyruvate and 30 % fetal calf serum (FCS) (Gibco, 10270-106). After the first passage the antibiotic concentration was reduced to 1%.

### Transfection of CRISPR/Cas9 plasmids

In total, 3×10^6^ cells were transfected when they reached 70-80% confluency. The two CRISPR/Cas9 plasmids (pX330-TRDCEX4#1+2) were co-transfected (at a final concentration of 5 μg/μl) into porcine fibroblasts by electroporation (Neon™ Transfection System, ThermoFisher Scientific) to test the efficacy of the plasmids to induce a 40bp deletion at the targeted locus (Figure X). Electroporation conditions were as follows: 1350 V, 20 mm, and two pulses. After lysis of transfected cells in cell lysis buffer followed by ethanol extraction, the DNA was analyzed using TRDC specific primer (TRDCEx4-2F: 5’-CTGGGTGTAAGTAGCAGCCT-3’ and TRDCEx4-2R: 5’-ACACGAGTTTTGAGTCTGGC-3’). The purified 499bp PCR product (10 ng/μl) (Invisorb® Fragment CleanUp – Startec) was Sanger sequenced to detect mutations at the target site. To produce TRDC-KO cell clones for serving as donor cells in SCNT, cells were trypsinized and subsequently diluted to a final concentration of 7-10 cells per well.

### In-Vitro-Fertilization and In-Vitro-Maturation

In-vitro-maturation of porcine oocytes was performed as previously described^39^. Briefly, porcine oocytes were collected from ovaries derived from slaughterhouse and matured for 40 hours in FLI medium. For in vitro fertilization, frozen boar semen from a fertile landrace boar was thawed for 30 seconds in a water bath (37 °C). The sperm motility was microscopically checked (Olympus, BH-2). After washing with Androhep® Plus (Minitube) and centrifugation for 6 minutes at 600 g, approx. 75 to 100 sperm per oocyte (depending on semen capacity) were used for fertilization (no sexed sperm were utilized for fertilization). After four hours of co-incubation, the fertilized oocytes were cultured in porcine-zygote-medium (PZM-3 medium).

### Intracytoplasmic Microinjection

Plasmids were prepared in 10 mM Tris-HCl pH 7.6 and 0.25 mM EDTA pH 8.0, and backfilled in glass injection capillaries and diluted to a final concentration of 2.5 ng/μl. Individual zygotes were fixed by suction to a holding pipette. The plasmids pX330-TRDCEX4#1 and pX330-TRDCEX4#2 were intracytoplasmically co-injected into IVF-produced zygotes derived from oocytes collected from slaughterhouse ovaries 20 hours after fertilization. To this end, approx. 10 pl plasmid solution was injected with a pressure of 600 hPa into IVF-produced zygotes (FemtoJet, Eppendorf). The injected zygotes were cultured in PZM-3 medium at 39 °C, 5 % CO_2_ and 5 % O_2_. At day 5, when embryos had reached the morula/blastocyst stage, 35 embryos were surgically transferred into each of the two recipients.

### Somatic cell nuclear transfer

SCNT was performed as previously described^40^. Fetal fibroblasts transfected with px330-TRDCEx4#1 and px330-TRDC-Ex4#2 targeting the exon 4 of the porcine TRDC gene were used as donor cells. In order to produce syngeneic wild type control pigs simultaneously, we chose cell clone D12 that gave two bands with a 50:50 ration on the PCR gel (Fig. 3) and cell clone H2 that gave an almost clean −40bp PCR band. In total, 90 H2-derived one-to two-cell embryos were surgically transferred into each of two hormonally synchronized German Landrace gilts (7 to 9-months old). For D12-derived embryos, 97 and 96 one-to two-cell embryos were surgically transferred into two hormonally synchronized German Landrace gilts mg/day/gilt Altrenogest (Regumate® 4mg/ml, MSD Germany) for 12 days, followed by an injection of 1,000 IU PMSG (pregnant mare serum gonadotropin, Pregmagon®, IDT Biologika) on day 13 and induction of ovulation by intramuscular injection of 500 IU hCG (human chorion gonadotropin, Ovogest®300, MSD Germany) 72 h after PMSG administration. The surgical embryo transfer was performed the day after the hCG administration.

### Genotyping of transfected cells and TRDC-KO pigs

Genomic DNA of the pigs was extracted from tail tips. Transfected cells and tail tips were lysed in cell lysis buffer (10%SDS, Proteinase K (20mg/ml), 10xPCR Buffer, aqua dest.) and purified by ethanol extraction. The DNA concentration was determined using the NanoDrop™ (Kikser-Biotech) system. For genotyping the pigs, polymerase chain reaction (PCR) was employed using specific primers (TRDCEx4-2F: 5’-CTGGGTGTAAGTAGCAGCCT-3’ and TRDCEx4-2R: 5’-ACACGAGTTTTGAGTCTGGC-3’) flanking a 499 bp segment of the TRDC gene (Fig. X). PCR amplification was performed in a total volume of 50 µl: 20 ng DNA, 0.6 µM reverse and forward primer, 1.5 mM MgCl_2_, 0.2 mM dNTPs and 1.25 U *Taq* Polymerase. Cycling conditions were as follows: 32 cycles with denaturation at 94°C for 30 sec, annealing at 60 °C for 45 sec, extension at 72°C for 30 sec and a final extension at 72°C for 5 minutes. The standard conditions for gel electrophoresis were set up to 80 V, 400 mA and 60 min using a 1 % agarose gel. The PCR-product was purified (Invisorb®Fragment CleanUp-Kit, Startec) and Sanger sequenced.

### Allele-specific sequencing of the TRDC gene of cloned piglets

The PCR product of the targeted region within exon 4 of the TRDC gene was subcloned into the pGEM-T Easy Vector system (Promega, Germany) in accordance with the manufacturer’s protocol and then transformed into XL10 bacteria. Bacteria were plated on Ampicillin containing Agar dishes (100 μg/ml) and allowed to grow for 16 hours. For each piglet, six colonies were picked and sequenced by using the T7prom primer.

### FACS analysis of TRDC-knockout blood and spleen cells

PBMC were isolated by density gradient centrifugation using Lympholyte®-Mammal Cell Separation Media (Cedarlane, Canada). Spleen cells were isolated by generation of a single cell suspension and erythrocyte lysis with ammonium chloride. For flow cytometry 1×10^5^ cells were incubated with primary antibodies followed by an incubation with secondary antibodies. The following primary antibodies were used anti-TCRγδ (PGBL22A, IgG1; PPT16, IgG2b), anti-CD2 (MSA4, IgG2a), anti-CD3 (PPT3, IgG1), anti-CD4 (74-12-4, IgG2b) and anti-CD8 (11/295/33, IgG2a). Detection of binding primary antibodies was performed using anti-mouse IgG1-PE, anti-mouse IgG2a-FITC, anti-mouse IgG2b-APC. All Incubations were done in acid buffer at 4°C and dead cells were excluded by adding propidium iodide (PI) (dilution 1:1000 in acid buffer) to the cells before measurement. Flow cytometry was performed with the MACSQuant Analyser and the “MACS Quantify” software.

### Transcriptome analysis

RNA from isolated PBMC was extracted using TRIzol® Reagent (Thermo Fisher Scientific). RNA from 4 TRDC-knockout pigs and 2 wild type pig were pooled, respectively. Pools were used to assess the quality of the RNA using an agilent bioanalyser. Labeled fragmented single-stranded cDNAs (ss-cDNA) were synthesized by using purified total RNA (100– 500Lng) as template following Affymetrix WT PLUS Labeling Assay protocols. Porcine Gene 1.1 ST Arrays (Affymetrix, Santa Clara, CA, USA) were hybridized to the biotinylated ss-cDNA targets. After 20□h of hybridization at 48□°C, arrays were washed by a fluidics station and then scanned by an imaging station in a GeneAtlas System (Affymetrix, Santa Clara, CA, USA). After scanning, the intensity data (CEL files) of Porcine Gene 1.1 ST arrays (Affymetrix) were extracted from the image data (DAT files) by the Affymetrix Command Console Software Version 1.4, and then normalized and analyzed by the Affymetrix Transcriptome Analysis Console (TAC) Software 4.0 for gene expression profiles and DEGs. The DEGs were selected by a cutoff of fold change >2.

### Autopsy, Histology and immune fluorescence

Full autopsy was performed on all pigs. Samples of the thymus, tonsil, spleen, femoral bone marrow, ileal Peyer’s patches, and lymph nodes (mandibular, tracheobronchial, cecal, popliteal) were immersion-fixed in neutral buffered, 10% formalin for at least 72 hours, and additional samples were snap frozen in liquid nitrogen.

Formalin-fixed tissues were trimmed, embedded in paraffin, cut in 2 μm sections and stained with hematoxylin and eosin (H&E) according to standardized procedures. For immunohistochemistry (IHC), consecutive slides were mounted on adhesive glass slides, dewaxed in xylene, followed by rehydration in descending graded alcohols. Endogenous peroxidase was quenched with 3% H_2_O_2_ in distilled water for 10 minutes at room temperature (RT). Heat induced antigen retrieval was performed in a decloaking chamber for 10 minutes at 110°C in 10mM citrate buffer (pH 6). Nonspecific antibody binding was blocked with 1:2 diluted goat normal serum in Tris-buffered saline (TBS) for 30 minutes at RT. A polyclonal rabbit anti-CD3 (#A0452, diluted 1:100 in TBS, Dako Agilent, Santa Clara, CA, USA) or rabbit anti-CD20 (#RB-9013-P1, diluted 1:200 in TBS, Thermo Fisher Scientific, Waltham, MA, USA) was applied over night at 4°C. A secondary biotinylated goat-anti rabbit antibody was used (#BA-1000, diluted 1:200 in TBS, Vector Laboratories, Burlingame, CA, USA) for 30 minutes at RT. The red-brown antigen labelling was developed by application of avidin-biotin-peroxidase complex (ABC) solution (Vectastain ABC Kit, #PK 6100, Vector Laboratories), followed by exposure to 3-amino-9-ethylcarbazole substrate (AEC, Dako, Agilent, Santa Clara, CA, USA). Sections were counter-stained with Mayer’s hematoxylin, dehydrated in ascending graded alcohols, cleared in xylene, and coverslipped. All slides were digitized using the Aperio CS2 slide scanner (Leica Biosystems Imaging Inc., CA, USA) and image files were generated using the NDP.view2 Software (Hamamatsu Photonics, Hamamatsu City, Japan). The establishment of immunofluorescence double-labelling using a polyclonal rabbit anti-CD3 and a monoclonal mouse anti-TCRγδ/PGBL22A antibody on snap frozen lymphoid tissues failed (not details shown). All examinations were performed by a board-certified pathologist (DiplECVP).

### Vaccination experiment

Five pigs at an age of 3 months were included in a vaccination study. Three of these animals had a γδ knockout phenotype (numbers 1314, 1318, 1319 (738-1, 737-4, 737-5) = group 1] and two represented syngenic wild type controls [group 2, numbers 1315, 1320 (737-1, 737-6)]. The two groups were kept in separate rooms. Vaccination was done after 14 days of acclimatization with a commercial life CSFV vaccine (Pestiffa CL, Boehringer Ingelheim Vetmedica, Ingelheim, Germany) according to the providers’ recommendation via the intramuscular route into the muscle brachiocephalicus. After vaccination, the animals were monitored daily. Rectal temperatures were recorded daily from −10 dpv to 24 dpv. Blood samples for serum production were taken on days 0 dpv and 21 dpv. Determination of neutralizing antibody levels was done as essentially as described before ^41,42^

## Declarations

### Ethics approval and consent to participate

Animal experiments were approved by the supervisory authorities (LAVES, AZ 33.19-42502-04-17/2532 and LALLF, 7221.3-2-042/17) and conducted in compliance with the German animal welfare law, the German guidelines for animal welfare and the EU Directive 2010/63/EU. All experiments were performed in accordance with relevant guidelines and regulations.

### Consent for publication

Not applicable.

### Availability of data and materials

Not applicable

## Authors’ Contributions

B.P., R.K. and G.M. conceived and designed the study and carried out data analysis as well as writing and revising the manuscript. B.P., A.F., P.H., R.B. and A.L.-H. established the TRDC-KO pigs, T.H.D., A.B. and R.G.U. performed experiments and data analysis.

## Acknowledgements

We thank Julia Sehl for the support with the autopsy and Gaby Stooß, Franziska Grenkowitz, Freja Pfirske, Gabriele Czerwinski and Maren Ziegler for excellent technical assistance.

